# Pan-cancer virtual spatial transcriptomics from routine histology with Phoenix

**DOI:** 10.64898/2026.04.25.720812

**Authors:** Manuel Tran, Rushin H. Gindra, Philipp Putze, Kang Senbai, Giovanni Palla, Tina Kos, Chiara Falcomatà, Chen Wang, Ruifeng (Ray) Guo, Melanie Boxberg, Luc M. Berclaz, Lars H. Lindner, Linda Bergmayr, Thomas Knösel, Philipp Jurmeister, Frederick Klauschen, Krisztian Homicsko, Raphael Gottardo, Markus Eckstein, Christian Matek, Andreas Mock, Fabian J. Theis, Dieter Saur, Tingying Peng

## Abstract

Spatial transcriptomics links gene expression to tissue architecture, providing a mechanistic view of cellular organization. Yet existing datasets cover few donors and miss the complexity of human disease. Experimental costs remain prohibitive, and large-scale profiling is impractically slow for population-level studies. Accurate computational methods are urgently needed. Predicting gene expression from standard histology, however, remains an open problem, as current approaches transfer poorly to unseen cohorts and diseases. Here, we present Phoenix, a (latent) flow matching generative model that infers pan-cancer spatially resolved single-cell gene expression with high accuracy. Phoenix analyzes treatment response in silico: Applied to 763 head and neck cancer patients, it identified three new spatial biomarkers that we validated across two cancers (breast cancer, n = 84; ovarian cancer, n = 157) and treatment regimens (platinum, trastuzumab). Phoenix generalizes beyond carcinomas: In a large sarcoma cohort (802 tissue microarray cores), it accurately predicted cell-type-specific signatures in held-out samples and captured chemotherapy-induced immune remodeling. Phoenix also extends across species: In a mouse model, it accurately predicted the expression of pancreatic cancer lineage markers and the mutant mKras^G12D allele in silico. In total, we evaluated Phoenix on over 10,000 patients. Our results establish virtual spatial transcriptomics as a scalable framework for studying tissue organization, therapeutic response, and disease mechanisms.

## Main

Spatial transcriptomics^1,2^ (ST) provides a quantitative readout of either whole transcriptome or targeted gene expression mapped to specific locations within the tissue. It has proven to be a powerful tool for understanding the activity and organization of cells in their native environment. Combined with histological staining and imaging, ST allows researchers to localize rare cells, characterize cell niches, visualize cell–cell interactions, trace signaling networks, uncover multiscale topologies, and identify biomarkers with greater precision.^3,4^

Technologies such as Visium^5^ map mRNA molecules in tissue neighborhoods (spots) at the whole transcriptome level and then sequence them ex vivo using NGS-based approaches. Since each spot is 55 µm in diameter, the typical readout measures expression from multiple cells simultaneously. To achieve single-cell resolution, platforms such as Xenium^6^ image mRNAs in situ and assign transcripts to individual cells through segmentation, but they are limited to a smaller number of genes. Subsequent methods, such as Visium HD^7^ and Xenium 5K^8^, achieve finer spatial resolution while capturing a greater number of genes.

Despite their successful applications in practice, current platforms share the same drawbacks: They require significant financial resources, typically costing thousands of dollars per sample, and take several days to complete. This makes ST inaccessible to many institutions, hinders large-scale efforts to study complex diseases such as cancer, and is a major barrier to clinical translation. Accurate computational methods are therefore urgently needed. However, predicting highly accurate gene expression from routine hematoxylin–eosin (H&E) histology images remains an open problem. Existing machine learning approaches fail to transfer robustly to new samples without extensive retraining because of batch effects, and perform reliably on less than hundred highly variable genes (HVGs). Consequently, prior work has neither demonstrated robust, population-scale performance nor convincingly tackled biomedical relevant questions across diverse cohorts, limiting the impact of deep learning in biological research and clinical applications.

We introduce Phoenix (Fig. 1), an end-to-end generative AI system that addresses these challenges. It improves upon the state-of-the-art by 35%-173% in terms of Spearman correlation (Fig. 1c). We achieved this through substantial advances in data filtering, model design, and compute scaling. In particular, we curated a dataset of 22.2 million cell-image & cell-expression pairs spanning 16 organ systems and 7 gene panels, and designed a family of deep neural networks based on our novel (latent) flow-matching architecture. We trained these models across multiple accelerators on the JURECA pre-exascale modular supercomputer, totaling more than 10,000 GPU hours – including our flagship human carcinoma model on the Xenium multi-tissue gene panel. At this scale, Phoenix works zero-shot (out-of-the-box, without fine-tuning) on new samples from diverse cohorts and donors (Fig. S1); extends to previously unseen tissues and organs; recapitulates key findings from published spatial studies; generates testable biological hypotheses; evaluates treatment responses in silico; screens clinical biomarkers at scale; and builds spatial atlases of over 10,000 patients. In sum, Phoenix establishes virtual spatial transcriptomics at scale.

**Fig. 1:**
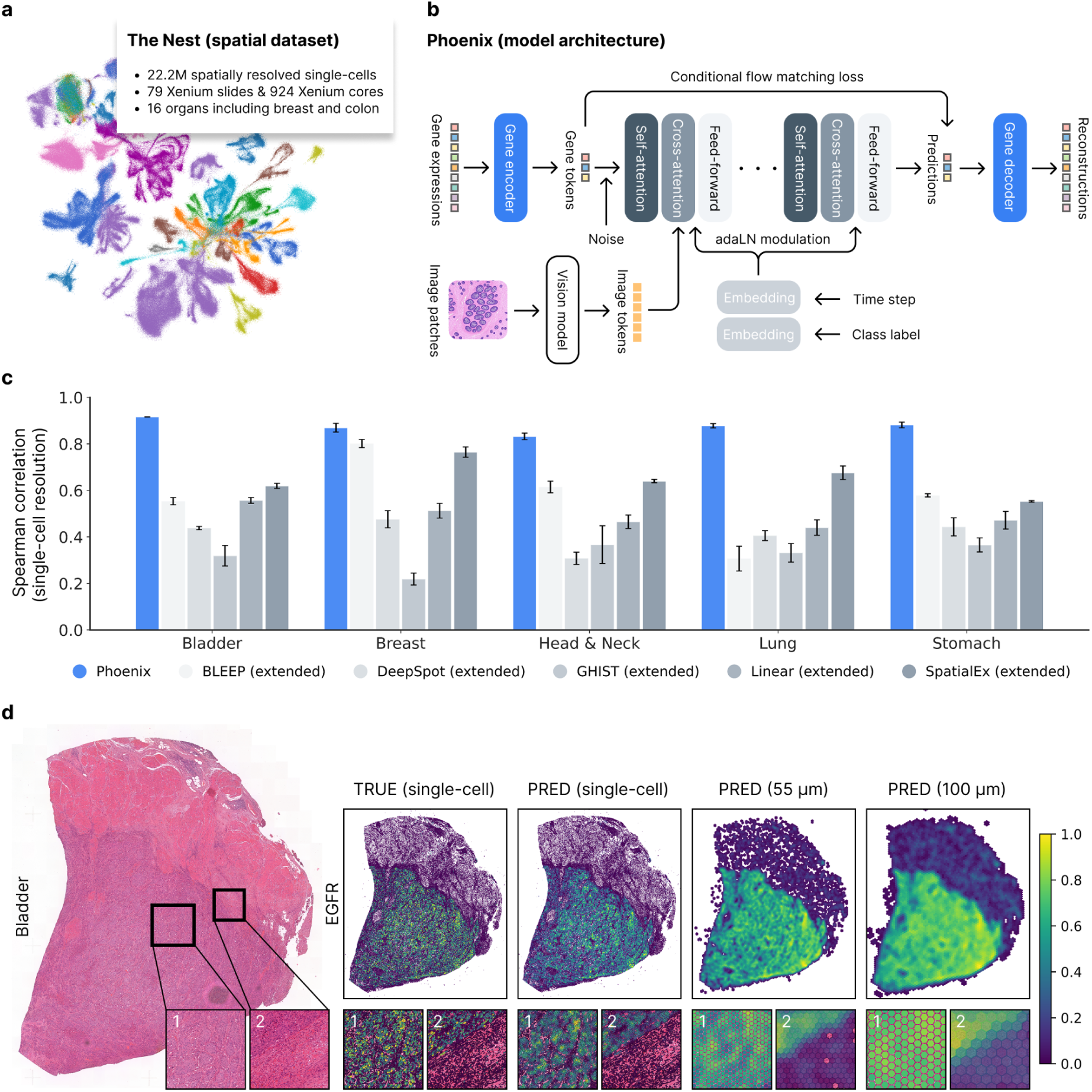
Phoenix accurately predicts gene expressions from histology images. **(a)** The Nest is a highly curated dataset of 79 Xenium whole slide images and 924 Xenium tissue microarray cores spanning 16 organ systems and 7 gene panels, totaling 22.2 million cell-image & cell-expression pairs. **(b)** Phoenix uses a (latent) flow matching model conditioned on pathology images to generate single-cell transcriptomics. **(c)** Zero-shot performance across organs (healthy, cancer) and Xenium gene panels (breast, lung, immuno, multi-tissue) on independent cohorts (see Fig. S1). Low and noisy expressions (counts ≤ 1) were binned to zero before computing Spearman correlations. **(d)** Examples of Phoenix predictions (EGFR) in a bladder cancer sample (TRUE = experimental ground-truth expression, PRED = out-of-domain model prediction); heatmaps are normalized and clipped at the 90th percentile.

### Scaling of data, model, and compute in spatial biology

We chose a data-driven approach to learn the relationship between morphological structures and gene expression profiles without relying on hand-crafted features (Fig. 1a,b). This makes the quality of the training data critical. Although Visium offers the largest collection of ST data, many samples suffer from severe batch effects (Fig. S1) – arising from low sensitivity^9^, increased sparsity^10^, lateral diffusion^11^, poor image quality^12^, and morphological distortion from fresh-frozen tissue^13^. We therefore trained exclusively on high-resolution imaging-based single-cell datasets of formalin-fixed, paraffin-embedded (FFPE) tissue sections, such as Xenium. Xenium is more sensitive (detecting a median of 12.8 times more reads than Visium^6^) and shows better concordance with spatial protein expression, suggesting that Visium especially suffers from platform-specific technical variation^10^. Most Xenium datasets also provide higher-resolution images (40× magnification, 0.25 µm/pixel) and use FFPE preparations that better preserve tissue morphology.

Our final dataset, The Nest, comprises 79 Xenium slides and 924 Xenium cores spanning 16 organ systems and 7 gene panels, totaling 22.2 million image–expression pairs from donors across North America, Europe, and Asia (Fig. 1a, Fig S1). It constitutes a large and diverse carcinoma cohort that combines a custom-curated subset of 10x Genomics’ public dataset (TENX) together with five external carcinoma cohorts (Methods; Fig. S1b): CHUV (Lausanne University Hospital), LMU (Ludwig Maximilian University of Munich), NCBI (Discovery Life Sciences), SNUH (Seoul National University Hospital), and UKER (University Hospital Erlangen). In addition, a second LMU cohort contributes human sarcoma samples, and the TUM (Technical University of Munich) cohort provides murine pancreatic cancer samples. Each panel captures a different level of biological context: the breast (280 genes), colon (322 genes), lung (289 genes), and skin (260 genes) panels focus on individual organs, while the immuno-oncology (380 genes) and multi-tissue & cancer (377 genes) panels are tissue-agnostic. Beyond single-cell resolution, we pooled transcripts into pseudo-spots of 55 µm and 100 µm to mimic Visium spots at two additional spatial scales (Fig. 1d, Fig. S1a). The 55 µm spots represent cellular niches, whereas the 100 µm spots reflect broader tissue domains. Both enable faster inference when single-cell resolution is unnecessary.

The data was then modeled using conditional flow matching^14^ (CFM). We adapted CFM for spatial biology by training a deep neural network^15^ called Phoenix. Phoenix consists of three components (Fig. 1b): an encoder that projects high-dimensional gene expressions into a lower-dimensional latent space; a flow model that generates latent vectors conditioned on image features; and a decoder that maps latent vectors back to gene expressions. The flow model is based on our modified transformer^16–19^, while the autoencoder (AE) is based on our custom redesign of the MLP-Mixer^20^. The AE is optional and can be replaced by the identity function (or potentially another gene foundation model like NicheFormer^21^), though at the expense of increased computational complexity and slower training and inference. A pathology foundation model (PFM)^22^ encodes the image into compact representations. This enables Phoenix to analyze gigapixel whole slide images (WSIs) in under one hour using off-the-shelf hardware, compared with 3–6 days on the Xenium In Situ platform.

### Generalization to unseen cohorts under batch effects

Predicting spatial expression from H&E images is only practical if models generalize zero-shot (out-of-the-box, without fine-tuning) on new samples from previously unseen donors, organs, institutions, scanners, and stainings. Proving this requires strict separation between training and evaluation. We therefore split the Nest at the cohort level: one training cohort and multiple held-out test cohorts. No donor, sample, or clinical center appears in both the training and test sets. For training, we chose the public TENX cohort (32 samples, 8.2 million cells), on which we optimized Phoenix, alongside state-of-the-art baselines (BLEEP^24^, DeepSpot^25^, GHIST^26^, Linear^27^, SpatialEx^28^). For testing, we evaluated all models on the external cohorts (Methods; Fig. S1): CHUV, LMU, NCBI, SNUH, and UKER. Together, these validation cohorts span five organs (bladder, breast, head and neck, lung, stomach), four gene panels (breast, lung, immuno, multi-tissue), and three continents (North America, Europe, Asia), forming a stringent and diverse benchmark for model generalization.

On this benchmark, Phoenix outperforms all baselines, improving correlation by 35–173% across cohorts, tissues, and panels (Fig. 1c, Fig. S2). Most notably, Phoenix works on unseen organs. Gastric samples were absent from the TENX training cohort (Fig. S1), making this an out-of-domain evaluation setting for Phoenix. Despite zero prior exposure, Phoenix predictions closely match Xenium measurements. On the other hand, competing methods collapse to the mean expression level (Fig. 2a): Neighboring cells or spots are assigned almost identical values. This eliminates spatial gradients, as confirmed by Geary’s C^29^ spatial autocorrelation (Fig. S2). Gradual, directional, and spatially organized expression changes are thus lost. Phoenix, however, does preserve biological structure at the tissue level. In a head and neck squamous cell carcinoma (HNSCC) lymph node metastasis (another tissue absent from training), Phoenix correctly predicts compartment-specific gene expression (Fig. 2b): CD3D in lymphoid tissue, EHF in more differentiated tumor regions surrounding keratinization, MET at the invasive tumor margin, and SFRP4 around the lymph node capsule at the interface with adjacent adipose and loose connective tissue. This accuracy extends to the cellular level, where label transfer recovers the same cell type annotations as experimental expression and outperforms all previous approaches (Fig. 2c,d).

**Fig. 2:**
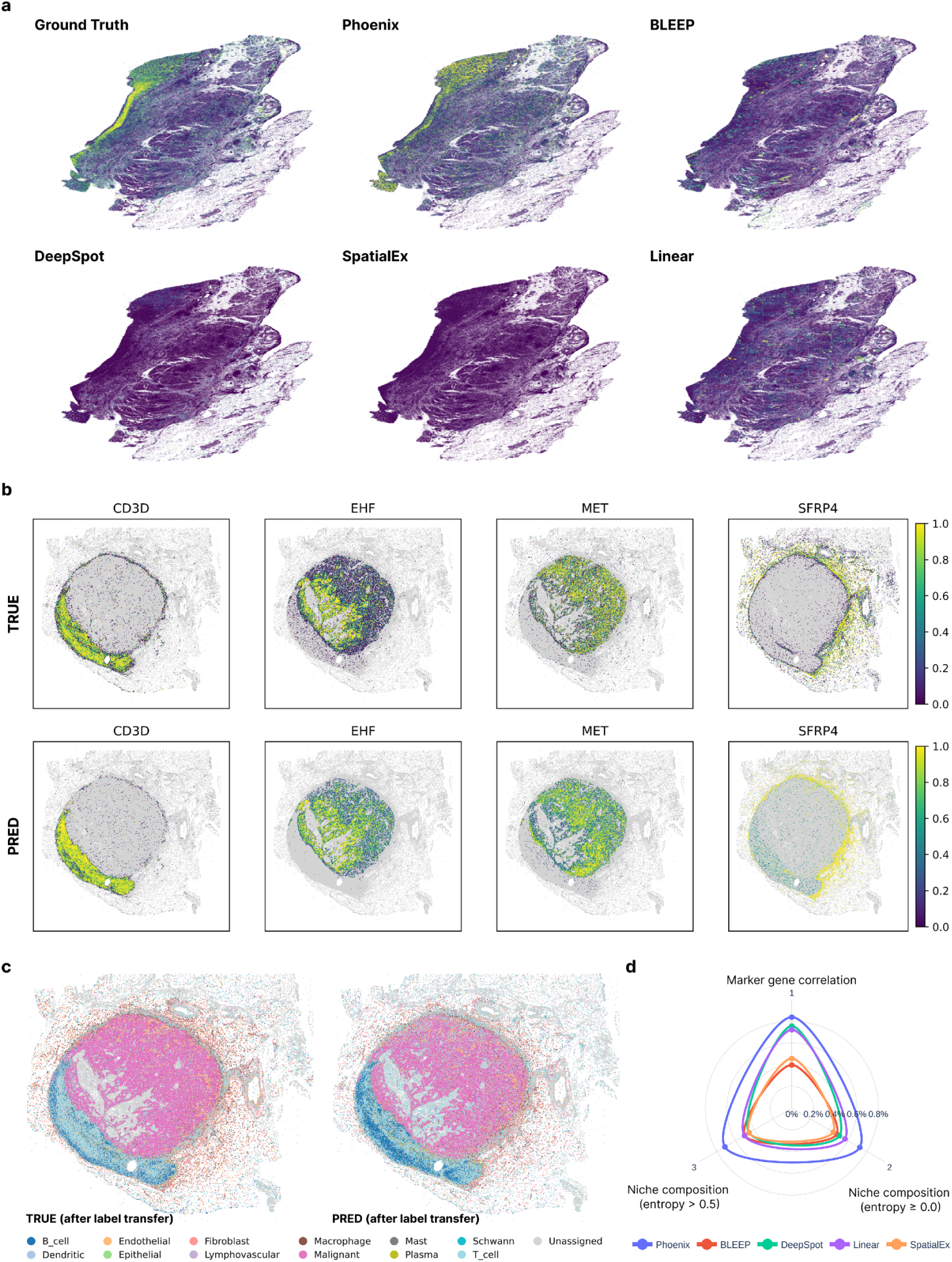
Phoenix resolves spatial architecture across unseen organs and tissues. **(a)** Local Geary’s C maps of the top-5 models visualizing the spatial gradient of highly variable genes in a gastric cancer — a tissue absent from the training set. **(b)** Phoenix predictions alongside the Xenium reference in a head and neck squamous cell carcinoma (HNSCC) lymph node metastasis (another tissue absent from training). **(c)** Cell types inferred by label transfer from an HNSCC single-cell reference atlas^23^ for the sample in (b). **(d)** Two accuracy metrics gauge the quality of cell type predictions: correlation between experimental and predicted signature genes, as well as overlap between experimental and predicted niche compositions (neighborhoods of ten cells). Shannon entropy measures cell neighborhood heterogeneity; niches with entropy >0.5 are classified as complex (highly mixed).

Having shown that Phoenix generalizes across independent cohorts, patients, organs, and tissues while surpassing baselines, we trained our flagship human carcinoma model on the union of CHUV, LMU, NCBI, SNUH, TENX, and UKER to maximize sample coverage and robustness. We reserve this model exclusively for downstream biological and clinical analyses where experimental ground-truth spatial expression is unavailable; no benchmark in this paper evaluates Phoenix on data seen during training. We then fine-tuned separate models for human sarcoma and mouse PDAC analyses (Fig. 6).

### Reliable hypothesis testing in biomedical experiments

A model becomes valuable when it helps biologists and clinicians generate and validate hypotheses that guide research. It should approximate biology well enough to inform discovery, not replace experiments. We therefore asked whether Phoenix could recover established biology and reveal clinically relevant spatial programs across diverse diseases. We first demonstrate this on breast cancer (BRCA) subtyping, a task central to prognosis and biology-informed treatment. Applied to breast data from *The Cancer Genome Atlas* (TCGA-BRCA), Phoenix recovered molecular subtypes with a mean AUROC of 0.826 ± 0.052 in five-fold cross-validation (Fig. 3a). Whereas prior work showed that snRNA-seq alone yields concordant subtypes^30^, Phoenix verified that virtual spatial transcriptomics recovers the same subtype structure at population-scale. Phoenix-derived expression profiles further resolved the immune landscape of the rare Claudin-low (CLOW) subtype, revealing reduced cytotoxic T cell activity and increased macrophage infiltration (Fig. 3b).^31,32^ Basal-like tumors, by contrast, harbor significantly more exhausted CD8⁺ T cells than luminal tumors (95% CI: 0.030–0.069, p = 0.001; Fig. 3c) – consistent with prior reports, but supported here on a 34-fold larger cohort (n = 1,125 vs. 33).

**Fig. 3:**
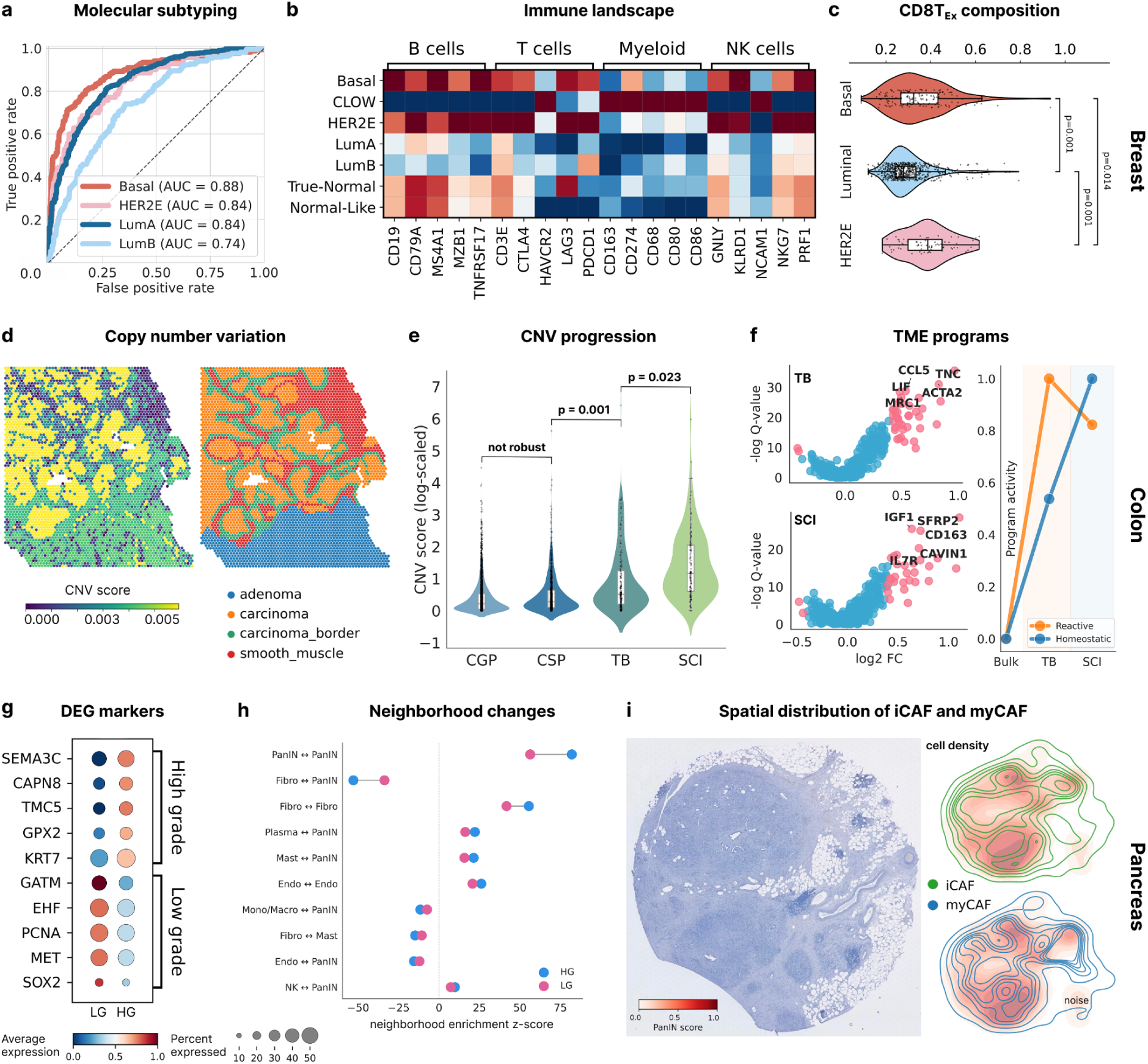
Phoenix discovers new biology and validates prior experiments. **(a)** In-silico spatial transcriptomics enables accurate molecular subtyping of breast cancer (BRCA) samples from TCGA using logistic regression (AUROC = 0.826 ± 0.052; five-fold cross-validation). (**b**) Immune marker expression varies across BRCA subtypes, revealing distinct tumor microenvironments (TME) along continuous subtype lineages (for example, LumA versus LumB and true normal versus normal-like). (**c**) Proportions of exhausted CD8⁺ T cells (Tex) relative to other T cells differ by BRCA subtype, with basal-like and HER2-enriched tumors showing significantly higher enrichment than luminal tumors (p = 0.001). **(d)** InferCNV applied to gene expression predictions recapitulates copy number variants (CNV) across distinct regions in colorectal cancer (CRC) tissue, closely matching pathology annotations. (**e**) High-resolution mapping powered by Phoenix identifies increasing genomic alterations across invasive growth patterns, from (adeno-)carcinoma with glandular growth pattern (CGP) and (adeno-)carcinoma with solid growth pattern (CSP) to tumor budding (TB; β ≥ 0.19, FDR < 0.002), and single-cell invasion (SCI; β ≥ 0.22, FDR < 0.025). **(f)** TB and SCI regions harbor TME niches distinct from the main tumor mass (FDR < 0.05), characterized by reactive (AIF1, LIF, TNC) and homeostatic (CD163, MRC1, SFRP2) immune programs. **(g)** The transition from low-grade (LG) to high-grade (HG) pancreatic intraepithelial neoplasia (PanIN) is marked by an increase of epithelial-mesenchymal transition (EMT) genes (e.g., SEMA3C) and a decrease of epithelial markers (e.g., EHF). **(h)** PanIN-HG exhibits more neoplastic cells and excludes fibroblasts, which encapsulate the lesion. **(i)** A sample with high-grade PanIN detected in the tissue, surrounded by myCAF fibroblasts with iCAF in the periphery.

We next used Phoenix to link histological architecture to regional evolutionary states in colorectal cancer (CRC). Applied to H&E images from Heiser et al.^33^, Phoenix captured the spatial mutational landscapes that shape CRC evolution (Fig. 3d). On a custom-annotated dataset^34^ spanning 32 morphology classes, Phoenix resolved the previously unclear link between pathology-defined structures and their underlying molecular alterations. We found that copy number variation (CNV) burden increases stepwise (FDR < 0.05) across invasive growth patterns (Fig. 3e), progressing from glandular (CGP) and solid (CSP) adenocarcinoma to areas of tumor budding (TB) and, finally, single-cell invasion (SCI) – mirroring the increasingly aggressive biology of CRC.^35^ Phoenix further distinguished the TME of TB and SCI regions (Fig. 3f). TB is enriched for AIF1, LIF, MRC1, and TNC (FDR < 0.05); and SCI for CD163 and SFRP2 (FDR < 0.05). Prior studies using flow cytometry, immunohistochemistry, protein assays, and survival analyses have linked these genes to poor prognosis and metastasis in colorectal cancer.^36–42^ Our these findings indicate that TB and SCI extend beyond EMT^43^, engaging distinct evolutionary programs of genomic instability and microenvironmental crosstalk (reactive vs. homeostatic TMEs).

Finally, we used Phoenix to resolve a preinvasive transition in the pancreas that remains difficult to profile experimentally at scale. Prior work identified gene programs that trace disease trajectories from pancreatitis through pancreatic intraepithelial neoplasia (PanIN) to pancreatic ductal adenocarcinoma (PDAC).^44^ Here, we focused on the poorly characterized transition from low-grade (PanIN-LG) to high-grade PanIN (PanIN-HG) in an independent cohort.^45^ We found that the progression from PanIN-LG to PanIN-HG involves the gain of invasive^46^ (SEMA3C) and tissue remodeling^47^ (CAPN8) programs, coupled with the loss of epithelial (EHF) and proliferative^48^ (MET) markers (Fig. 3g, Fig. S3).^49–52^ Together, these shifts reshape tissue architecture (Fig. 3h). Phoenix further partitioned the stroma into spatially distinct fibroblast niches: myofibroblastic CAFs (myCAFs) encapsulate PanIN clusters, while inflammatory CAFs (iCAFs) occupy segregated niches in the adjacent stroma (Fig. 3i)^53^.

### A pan-cancer spatial atlas of over 9,000 patients

Having established Phoenix through benchmark studies, biological validation, and clinical endpoints, we applied our model to patient cohorts at a scale previously inaccessible to spatial transcriptomics. Thus, we inferred spatially resolved gene expression across the entire TCGA (The Cancer Genome Atlas) project, spanning 9,544 donors, and combined these predictions with matched H&E-stained whole slide images to build a multimodal spatial atlas of human cancer (Fig. 4a,b). At three resolutions (100 µm, 55 µm, and single-cell), the atlas charts spatial hierarchies from broad tissue domains to local cellular niches. Phoenix pseudobulk profiles showed good concordance with TCGA bulk RNA-seq across 373 shared genes (Spearman ρ = 0.66 for mean expression, ρ = 0.53 for cross-patient variance), confirming the validity of the inferred expression values.

**Fig. 4:**
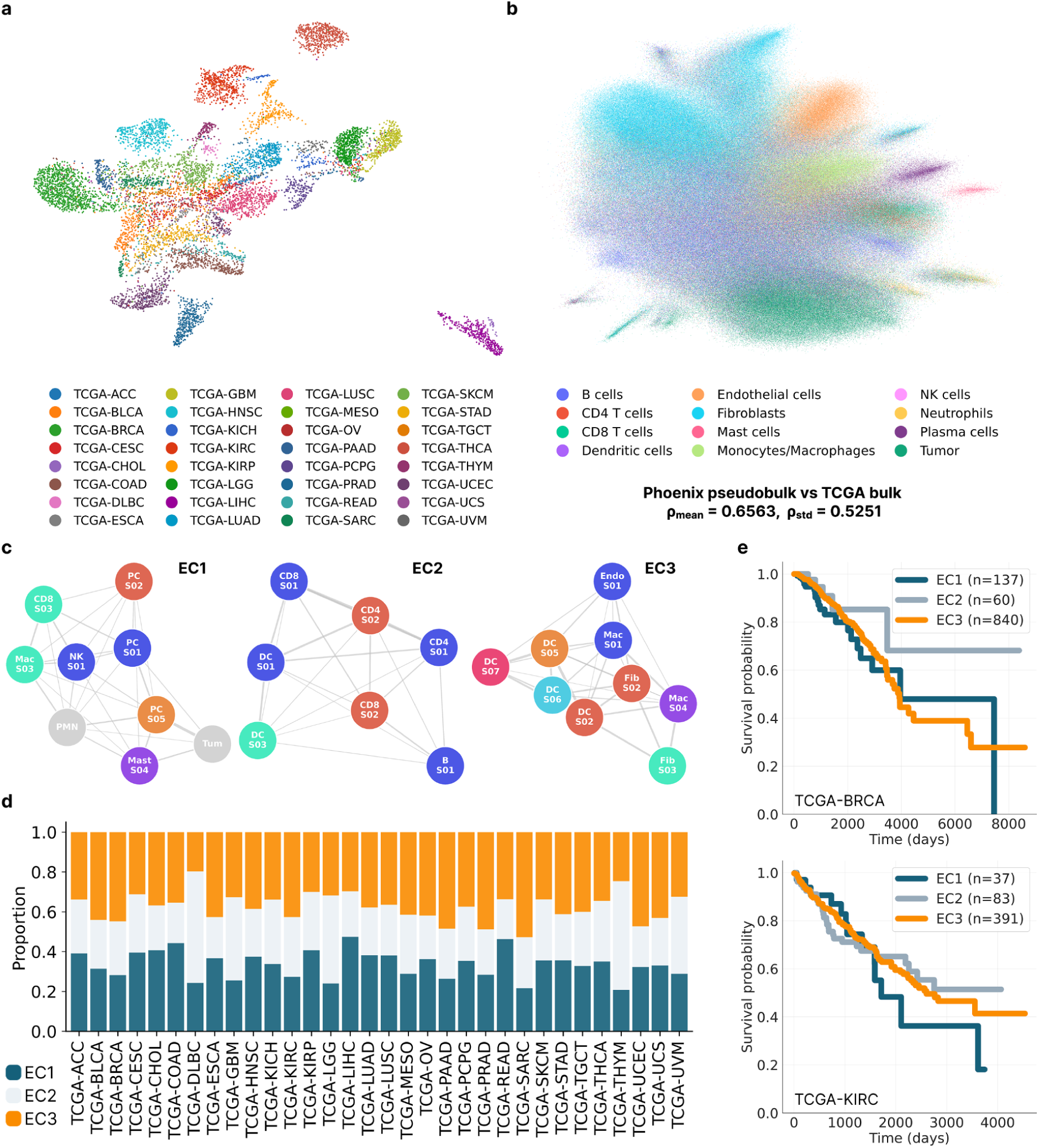
Phoenix scales to population-level datasets and delivers new insights. **(a)** A spatially resolved single-cell atlas of 9,544 patients derived from The Cancer Genome Atlas (TCGA) using Phoenix (one dot represents one patient). **(b)** Annotations reveal that virtual expression profiles cluster by cell type rather than tissue type (one dot represents one cell). Overall, Phoenix expression profiles shows good concordance with TCGA bulk RNA-seq across 373 shared genes (ρ_mean_ = 0.66, ρ_std_ = 0.53). **(c)** Ecotyper^54^ defines ecotypes as the co-occurrence of deconvoluted states (see Fig. S4 for more details); we extend this framework by incorporating spatial proximity between cells to identify niches. B: B cells, CD4: CD4 T cells, CD8: CD8 T cells, DC: dendritic cells, Endo: endothelial cells, Fib: fibroblasts, Mast: mast cells, Mac: monocytes/macrophages, NK: NK cells, PMN: neutrophils, PC: plasma cells, Tum: epithelial cells including tumor cells. **(d)** Distribution of three ecotypes (EC1, EC2, and EC3) across cancer types. **(e)** Kaplan–Meier survival estimates for breast (BRCA; top) and kidney (KIRC; bottom) carcinoma, stratified by dominant ecotype (winner-takes-all assignment based on the highest per-patient proportion). Group sizes therefore differ from the continuous proportions in (d) because patients with mixed ecotype profiles are classified by their majority component.

At the population scale, a topological principle of tumor organization emerges: Recurrent transcriptional states (denoted S0X, e.g., S01; see Fig. S4) assemble into conserved multicellular communities across carcinomas. Mapping their spatial proximity revealed three distinct ecotypes^54,55^ that shape the tumor microenvironment (Fig. 4c,d) – EC1 (immune dysfunction), EC2 (immune competence), and EC3 (immune exclusion). EC1 is enriched for pro-inflammatory macrophages, mast cells, neutrophils, NK cells, plasma cells, and exhausted CD8+ T cells, consistent with a hyperactive yet ineffective immune response. EC2 forms a coordinated immune hub built around cDC1-mediated antigen presentation and productive T-cell–B-cell interactions, matching a model in which mature cDC1s prime naïve CD4+ T cells, which then engage classical B cells and license cDC1s to cross-prime naïve CD8+ T cells.^56^ EC3 defines a remodeling compartment dominated by endothelial cells, fibroblasts, monocytes, and M2-like macrophages, compatible with matrix remodeling and tissue repair. Phoenix hence not only recovers cell types or genes, it also reveals conserved higher-order ecosystems.

Survival analysis across 32 cancer types showed that these ecosystems are also clinically informative. On average, the qualified immune response of EC2 confers strong survival benefits and defines the most protective baseline. By contrast, the exhausted state of EC1 markedly increases mortality risk (HR = 2.11, CI = 1.22–3.63, p = 0.007), and EC2’s advantage persists even after adjusting for tumor cellularity (HR = 0.87, CI = 0.78–0.97, p = 0.016). Stromal remodeling in EC3, though less detrimental than EC1, trends toward poorer prognosis than EC2 (HR = 1.59, CI = 0.95–2.68, p = 0.08). The ordering represents a general trend across TCGA rather than a uniform effect in every cancer, exemplified here by breast invasive carcinoma (Fig. 4e, upper panel). In some tumor types, notably kidney renal clear cell carcinoma, EC2 and EC3 confer comparable outcomes (Fig. 4e, lower panel). Overall, these findings suggest a refinement of Dvorak’s classic axiom^57^: Tumors are wounds that fail to heal, but they fail in distinct ways. Some remain locked in chronic inflammatory dysfunction (EC1), others progress toward aberrant fibrotic scarring (EC3), and favorable prognosis tracks with preserved immune competence (EC2).

### In silico modeling of treatment response mechanisms

A key question is whether Phoenix can leverage routine histology to model treatment response and uncover transferable biomarkers of therapeutic benefit across cancers and regimens. In an independent, public cohort of ovarian cancer patients treated with bevacizumab (n = 78),^58^ Phoenix recovered response-associated tissue states consistent with anti-angiogenic biology.^59^ We found that responders tend to have more immune-infiltrated / immune-deserted phenotypes (Fig. 5a, p = 0.0095) and more proliferative tumors (Fig. 5b, p = 0.013), compatible with prior reports.^60^ In addition, the proliferative subtype is overrepresented among high-risk responders (FIGO III–IV, p = 0.023), who derive greater benefit from treatment.^61^ It is known that proliferative tumors are oxygen-deprived, activating hypoxic stress signals and driving vascular angiogenesis through VEGF.^62^ VEGF inhibition, in turn, can engage alternative pathways, including TGF-β signaling.^63,64^ Notably, Phoenix identified BAMBI as upregulated in the TME of responders (FDR = 0.051–0.117), which aligns with attenuation of compensatory TGF-β signaling during VEGF blockade.^65^ Together, these findings show that Phoenix moves beyond response stratification to generate mechanistic hypotheses directly from archived tissue sections.

**Fig. 5:**
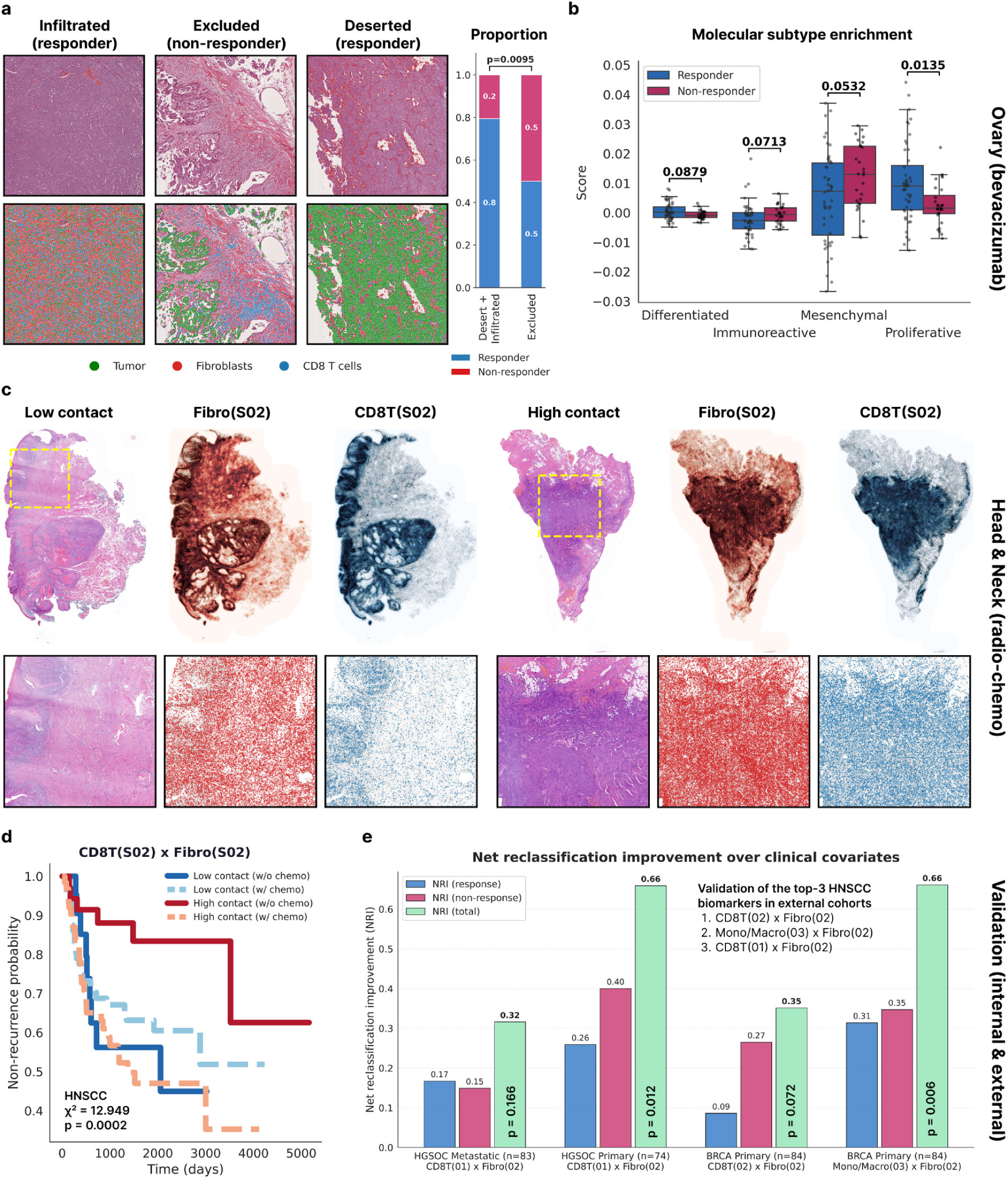
Phoenix models drug response mechanisms in silico across cancers. **(a-b)** Ovarian cancer bevacizumab therapy response (n = 78). **(a)** Immune landscape in responders versus non-responders. **(b)** Molecular subtype enrichment in responders versus non-responders. **(c-e)** HNSCC radiochemotherapy response (n = 763). **(c)** Spatial organization of Fibro(S02) and CD8T(S02) in low-contact compared to high-contact groups. **(d)** Kaplan-Meier recurrence estimates in HNSCC stratified by platinum chemotherapy and spatial co-occurence of CD8T(S02) × Fibro(S02). **(e)** Continuous Net Reclassification Improvement (NRI) of the top-3 HNSCC predictive biomarkers across cancer types (HGSOC, n = 157; BRCA, n = 84) and treatment regimens (platinum, trastuzumab).

Radiotherapy resistance severely limits survival in head and neck squamous cell carcinoma (HNSCC). We analyzed 763 patients from a clinical cohort in Erlangen, applying Phoenix to generate spatial profiles across four complementary axes: cell abundance, cell neighbors, cell niches, and cell location (Fig. S5). Patients were then stratified by whether they received adjuvant radiotherapy after surgery or surgery alone, as determined by disease stage. In the adjuvant-treated group, recurrent tumors show marked depletion of myeloid dendritic cells and T cells, both inside and outside the tumor area, with the few remaining T cells confined to fibroblast- and monocyte/macrophage-dominated niches (Fig. S5, p < 0.05). Recurrent tumors in the surgery-only group, on the other hand, selectively lose neutrophils (Fig. S5, p < 0.05). These distinct spatial signatures demonstrate that Phoenix resolves therapy-specific modes of immune failure rather than generic recurrence states.

A subset of patients also received concurrent chemotherapy on top of radiotherapy after surgery (adjuvant radiochemotherapy). We predicted drug benefit using a two-stage approach within repeated five-fold nested cross-validation. To explicitly identify *predictive* rather than purely *prognostic* markers, our first stage screened features using a formal treatment-by-biomarker interaction test. Specifically, we used a Likelihood Ratio Test (LRT) comparing Cox Proportional Hazards models with and without the interaction term, while strictly adjusting for a baseline clinical prognostic index (age, pN stage, pT stage, sex, smoking status) to prevent confounding by indication. Significant features were then re-fitted in a multivariate Ridge-Cox regression.^66–77^ Neighborhood information improves the C-index by 0.02 on average (95% CI: 0.01–0.03, p = 0.005), showing that cell–cell context contributes predictive information beyond cell abundance alone. In 89% of outer folds, spatial co-occurrence (high vs. low) between activated CD8⁺ T cells (S02) and resident cancer-associated fibroblasts (S02) emerged as the most robust predictive spatial biomarker for chemotherapy response (Fig. 5c).

The hazard ratio remained stable across folds (HR: 3.00–4.04, p < 0.01), and permutation of treatment labels confirmed the robustness of this interaction (χ² = 12.94, p < 0.001). Importantly, our interaction model adjusts for age, pN stage, pT stage, sex, and smoking status; and all resections were collected before systemic therapy. We next stratified patients by CD8T(S02) × Fibro(S02) interaction (Fig. 5d), revealing a clear clinical dichotomy: In the low-contact group, chemotherapy confers no detectable survival benefit (HR = 0.85, 95% CI: 0.39–1.82, p = 0.681), whereas the high-contact group experiences significantly worse outcomes (HR = 4.46, 95% CI: 1.80–11.09, p = 0.001). The second and third most consistently ranked biomarkers follow a similar pattern (Fig. S5). Critically, cell abundance alone does not predict outcome — only their interaction is informative.

Finally, we reasoned that spatial biomarkers capturing fundamental immune-stromal interactions should generalize across diseases and therapies. Across two independent cohorts of breast cancer treated with trastuzumab^78^ (BRCA, n = 84) and ovarian cancer treated with platinum^79^ (HGSOC, n = 157), the three top-ranked HNSCC biomarkers improve risk stratification, quantified by the continuous Net Reclassification Index (NRI; Fig. 5e). In BRCA, the first-ranked biomarker CD8T(S02) × Fibro(S02) shows a trend (NRI = 0.35, 95% CI: −0.04–0.75, p = 0.072), while the second-ranked biomarker Mono/Macro(S03) × Fibro(S02) has an even stronger effect (NRI = 0.66, 95% CI: 0.22–1.06, p = 0.006). In HGSOC, where CD8T(S01) rather than CD8T(S02) dominates the CD8 compartment,^54^ the third-ranked HNSCC biomarker CD8T(S01) × Fibro(S02) improves risk stratification in primary non-metastatic tumors (NRI = 0.65, 95% CI: 0.18–1.12, p = 0.012) but not in metastatic ones (NRI = 0.31, 95% CI: −0.12–0.77, p = 0.166).

Collectively, these results uncover spatial co-occurrence between immune and stromal cells that predict how patients respond across diverse tumor types and treatment regimens, all from archived pathology material. Phoenix offers a platform to model these therapeutic mechanisms i*n silico* at a population scale.

### Extension to mesenchymal cancers and mouse models

To rigorously evaluate transferability across lineages, species, and therapeutic perturbations, we first applied Phoenix to sarcomas. Sarcomas are rare cancers of mesenchymal origin, accounting for approximately 1% of all adult cancer diagnoses. Compared to carcinomas, they remain largely unexplored by spatial methods. We profiled 802 tissue microarray (TMA) cores spanning 15 sarcoma entities with Xenium In Situ. Because Phoenix was originally trained on carcinomas, we fine-tuned the model on our sarcoma data and evaluated it on 197 held-out cores from independent patients. Despite this lineage shift, Phoenix accurately reconstructed spatial gene expression across all 197 test cores, reaching a median micro-bulk Spearman correlation of 0.734 for the top 50 genes and 0.436 for all 377 genes. Notably, Phoenix recovered cell-type programs across the tumor microenvironment (Fig. 6a),^80^ with correlations exceeding 0.5 for 9 of 11 cell types (Fig. 6b), including monocytes/macrophages (rho = 0.813), CD4 T cells (rho = 0.748), endothelial cells (rho = 0.723), and CD8 T cells (rho = 0.702).

**Fig. 6:**
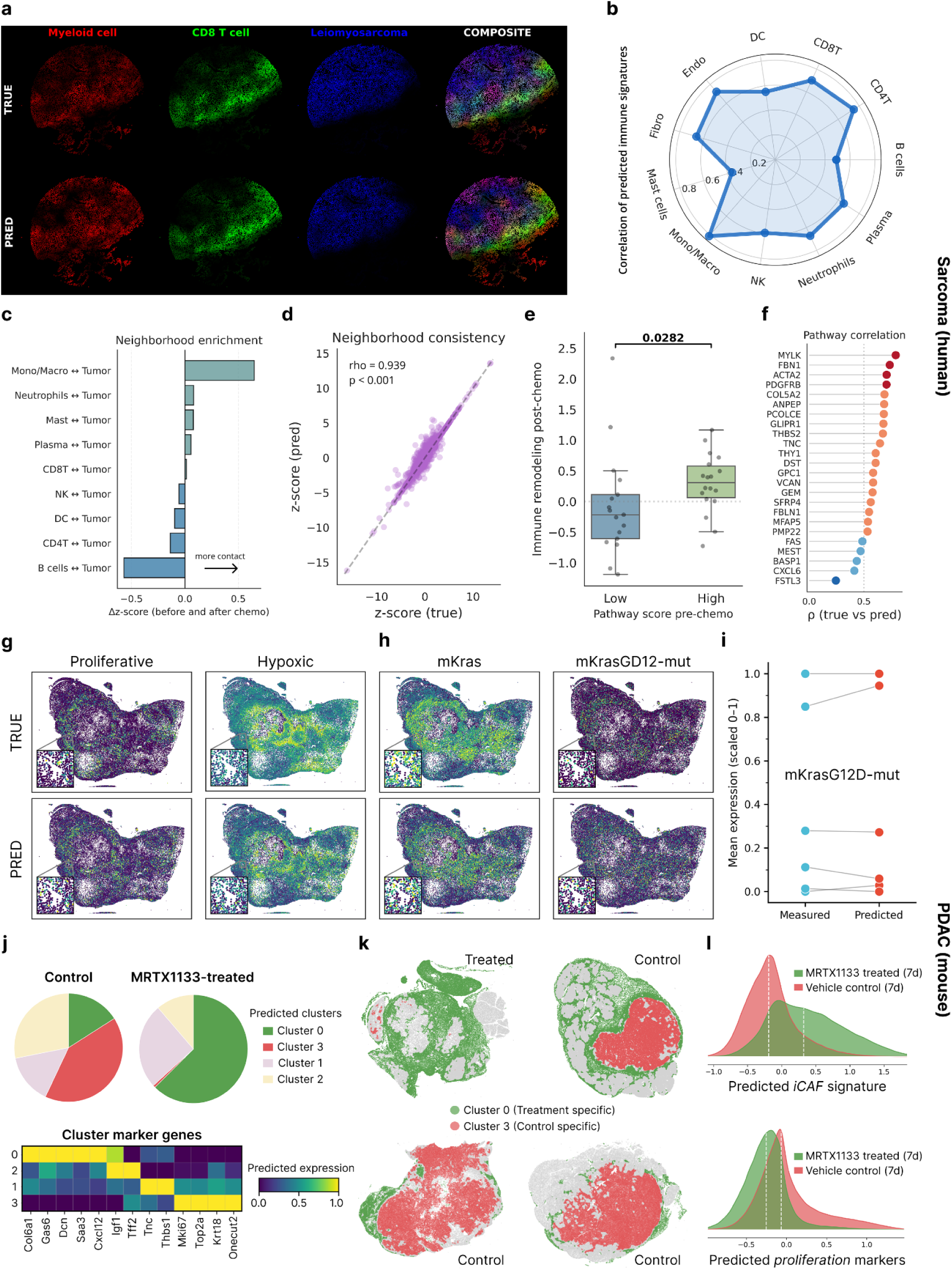
Phoenix screens clinically actionable biomarkers in silico. **(a-f)** Human sarcoma. **(a)** Molecular pseudocolor maps of myeloid markers (AIF1, CD14, CD68; red), CD8 T cell markers (CD3D, CD3E, CD8A; green), leiomyosarcoma markers defined by genes of smooth muscle differentiation (ACTA2, ACTG2, CNN1, DES, MYH11, MYLK; blue), and their composite in a TMA core (higher intensity = higher expression). **(b)** Spearman correlations of predicted immune cell markers on held-out test cores. **(c)** Neighborhood enrichment analysis before and after chemotherapy on experimental measurements. **(d)** Correlation of predicted spatial neighbors on held-out test cores. **(e)** M5930 GSEA pathway expression score before chemotherapy correlates with immune-remodeling after chemotherapy on experimental measurements. **(f)** Correlation of predicted pathway genes on held-out test cores. **(g-j)** Mouse PDAC. **(g)** Experimental and predicted spatial expression of mPDAC subtype markers: proliferative (Mki67, Top2a), hypoxic (Hif1a, Vegfa). **(h)** Experimental and predicted spatial expression of mKras and mKrasG12D-mut. **(i)** Experimental and predicted mean expression of mKras and mKrasG12D-mut across six mPDAC specimens. **(j)** Relative abundance of CellCharter clusters in vehicle control and MRTX1133-treated samples (top). Matrixplot displaying top marker genes per cluster (bottom). **(k)** Spatial distribution of Cluster 0 (treatment-enriched) and Cluster 3 (control-specific) across test samples. **(l)** Comparison of iCAF and proliferative gene signatures between MRTX1133-treated and vehicle-treated control samples after 7 days of therapy. iCAF scores are increased, while proliferation scores are decreased upon treatment (Wilcoxon, p < 1e-16).

In a matched subset of 51 sarcomas sampled before and after treatment, chemotherapy remodels the spatial immune architecture (Fig. 6c). On the held-out test cores, Phoenix predicted these spatial neighborhoods with near-perfect accuracy (Fig. 6d; rho = 0.939, p < 0.001), supporting its use for in silico analysis of treatment-induced remodeling. To identify pre-treatment features linked to subsequent immune remodeling, we screened gene sets across all 51 matched cores. GSEA (*Gene Set Enrichment Analysis*) pathway M5930, which includes ACTA2 and ANPEP, tracks with greater post-chemotherapy shifts in immune composition, including B cell depletion and enrichment of myeloid-dominated cancer cell neighborhoods (Fig. 6e, p = 0.028). PDGFRB shows the strongest individual association with immune remodeling (Fig. S6, FDR = 0.034), although its own expression does not change upon chemotherapy (paired Wilcoxon, p = 0.46). Overall, Phoenix predicted the expression of PDGFRB and the remaining M5930 pathway genes in held-out test cores with a mean correlation of 0.683 (Fig. 6f).

Phoenix is not tied to any single species or gene panel. Leveraging its foundation model design, we fine-tuned Phoenix on an in-house Xenium In Situ dataset from an established mouse model of pancreatic ductal adenocarcinoma (mPDAC), with six samples used for training and nine for testing. To capture the continuous spectrum of PDAC progression (acinar, epithelial, proliferative, hypoxic, epithelial-mesenchymal transition, and mesenchymal),^81,82^ we derived a 6-dimensional biomarker score representing relative subtype and cell-state identity from transcriptomic signatures. Predicted subtype distributions closely matched the ground truth (Fig. 6g, Fig. S6), with 90.7 ± 4.2% overlap at the sample level (JSD = 0.08 ± 0.05) and 79.8 ± 4.2% at the niche level (JSD = 0.17 ± 0.04).

Beyond cell-state identity, Phoenix accurately quantified total mKras and mutant mKras^G12D expression in situ (rho = 0.76), demonstrating its potential for patient stratification (Fig. 6h,i). In particular, these expression profiles revealed transcriptional remodeling in mPDAC following mKras^G12D inhibition with MRTX1133. Using CellCharter, we identified four distinct spatial domains, including a proliferative tumor cell niche (Cluster 3) dominant in untreated samples and an inflammatory CAF (iCAF) niche (Cluster 0) enriched in MRTX1133-treated tumors (Fig. 6j,k). Gene signature analysis confirmed a significant ecological shift from proliferative tumor-cell profiles to iCAF-enriched niches upon treatment, consistent with microenvironmental remodeling (Fig. 6l).^83^

Together, these findings extend Phoenix beyond human carcinomas to mesenchymal cancers and mouse models, and show that it can resolve lineage state, mutant allele expression, as well as therapy-induced ecosystem changes across lineage and species.

## Discussion

Phoenix shows that predictive methods for spatial transcriptomics require the joint scaling of data, model, and compute. Doing so is non-trivial, as existing approaches have not achieved robust-scaling across diverse cohorts, organs, and tissues. We therefore curated a high-quality Xenium dataset with 22.2 million single cells and designed a powerful generative architecture with 1.2 billion parameters. The resulting flow model correlates morphological features encoded in pathology images and translates them into transcriptional space without any imposition of handcrafted features. For example, Phoenix transfers molecular lineages across tissues, despite never being trained on class labels.

Phoenix generalizes across unseen donors, organs, and treatments — a capability no other method has demonstrated. This enables screening of thousands of patients in pathology archives at a fraction of the cost and time, opening the door to population-level spatial transcriptomics. We demonstrated this through hypothesis generation and validation, uncovering novel biology and actionable biomarkers including mesenchymal cancers and in vivo mouse studies. Notably, we discovered that specific configurations of the immune landscape emerge as a hallmark of treatment response across cancer types and therapies.

Although we used Phoenix in an international, multi-institutional effort to profile and screen over 10,000 specimens from public and private cohorts, only a consortium-level endeavor can unlock the full potential of hidden pathology archives with richer clinical endpoints. To this end, we release both model code and weights, enabling the community to explore spatial biology at unprecedented scale. This opens the possibility of profiling entire populations, bridging discovery science and clinical practice. As access to graphics processing units (GPUs) expands in clinical settings, we envision that Phoenix and its successors will become more widely deployable across medical centers worldwide.

A limitation of Phoenix is the restricted gene panel imposed by Xenium In Situ. Even though Phoenix can be readily fine-tuned on new panels, such as our mouse mPDAC panel, broader gene coverage would be welcome. Our research indicates that the bottleneck is primarily caused by poor data quality, not flawed model design. As a demonstration, we trained Phoenix on a diverse cohort of private and public Xenium 5K Prime data – it failed to generalize to new samples. Even subsetting the large panel to a smaller panel (e.g., breast with 280 genes), the model does not generalize (Fig. S7). After ruling out tissue type and image quality as potential confounders, we concluded that lower sensitivity is the main shortcoming of Xenium 5K Prime. Phoenix is trained and validated exclusively on Xenium in Situ data, and we acknowledge that its transferability to other in situ platforms (e.g., MERFISH, CosMx, or seqFISH+) remains untested. While the image encoder is platform-agnostic, gene expression distributions are platform-specific, which poses a potential challenge to cross-platform generalizability. Furthermore, cell type annotation in spatial transcriptomics remains constrained by panel composition, marker specificity, platform sensitivity, and methodological choice — limitations our analyses inherit. Nevertheless, the resulting spatial distributions and downstream characterizations are robust: They align with expected tissue organization and the canonical biological literature. By developing an accurate and scalable spatial transcriptomics inference engine, coupled with the growing availability of digital pathology archives, Phoenix can accelerate the adoption of computational biology that keeps pace with the sequencing revolution. We envision that Phoenix will become an essential tool of modern biomedicine and pathology.

## Methods

### Donor cohorts

The CHUV cohort comprises twelve breast cancer and five lung cancer samples from the Department of Clinical Oncology at Lausanne University Hospital, profiled with the breast and lung panels, respectively.^84^ Five additional lung cancer samples were profiled with a custom immuno-oncology panel. The LMU cohort, from the Institute of Pathology at the University of Munich, contributes one head and neck squamous cell carcinoma lymph node metastasis sample, 178 head and neck cancer cores, and 802 sarcoma cores, all profiled with the multi-tissue panel. The NCBI cohort includes three breast cancer samples from Discovery Life Sciences, profiled with the breast panel. The SNUH cohort contains two gastric cancer samples and one normal gastric sample from Seoul National University Hospital, and the UKER cohort adds three bladder cancer samples from the Institute of Pathology at University Hospital Erlangen; both were profiled with the multi-tissue panel. The TUM cohort, from the Center for Translational Cancer Research at the Technical University of Munich, provides 15 samples, profiled with the mPDAC panel.

The TENX cohort, obtained from publicly available datasets released by 10x Genomics, comprises 32 samples spanning diverse tissues and disease states. Six breast cancer samples (invasive ductal and lobular carcinoma) were profiled with the breast panel, and two colon samples (cancer and non-diseased) with the colon panel. One lung cancer sample was profiled with the lung panel, and two melanoma samples with the skin panel. Five samples covering brain, colon, lung, ovary, and pancreas cancers were profiled with the immuno-oncology panel. The remaining sixteen samples – spanning bone, bone marrow (including acute lymphoid leukemia), heart, kidney, liver, lung, pancreas, skin, and tonsil, in both diseased and non-diseased states – were profiled with the multi-tissue panel.

### Dataset construction

The color, quality, resolution, staining protocols, scanner settings, and tissue preparation of histopathology images vary widely, introducing variability that hinders robust analysis.^85–87^ We therefore defined high-quality images as those acquired at native 40× magnification (0.25 ± 25% µm per pixel) and formalin-fixed, paraffin-embedded (FFPE) tissue preparation.^12,13^ Applying these criteria to public and private collections yielded 79 Xenium slides and 924 Xenium cores spanning 16 organ systems and 7 gene panels. We excluded all Visium data due to lower sensitivity, lateral diffusion, and other artifacts.^6,9,11^

### Multimodal alignment

Multimodal alignment was performed by registering post-Xenium stained FFPE tissue sections onto their corresponding DAPI morphology maps. Transcript data exported from the Xenium platform were first filtered to remove negative controls and unassigned or deprecated codewords. All samples within The Nest were subsequently realigned using 10x Genomics Xenium Explorer to minimize human error; 30 anatomical landmarks per sample were identified as optimal for achieving robust one-to-one morphology-to-transcriptome correspondence. Aligned samples were serialized as Zarr objects within the SpatialData framework to provide a standardized, format-consistent representation suitable for downstream machine learning training. Following quality filtering, transcripts were aggregated into pseudo-spots at two spatial resolutions: 55 µm and 100 µm to simulate Visium-scale capture zones reflecting cellular niches and broader tissue domains, respectively. Spot aggregation was implemented using a hexagonal tiling scheme with 6% inter-hexagon overlap, wherein all transcript counts falling within a given hexagon were summed to generate the spot-level expression vector. Hexagonal rather than circular spot geometries were employed to ensure complete partitioning of the tissue area and eliminate transcript loss in inter-spot gaps. Spots with total gene count < 1 were filtered out.

### Image preprocessing

We tessellated each whole-slide image at 20× (0.50 µm/pixel) or 40× (0.25 µm/pixel) magnification into 224 × 224-pixel patches centered on a cell or spot, then extracted embeddings using one of four state-of-the-art pathology foundation models: HO-Mini^88^, H-Optimus-1^89^, UNI2-h^90^, and Virchow2^91^. All four are DINOv2-pretrained^92^ Vision Transformers^93^ with 87M, 1.135B, 681M, and 631M parameters, respectively. Each model produces a 16 × 16 feature grid, where each token represents a 14 × 14-pixel region. For the Flow Transformer, we passed all tokens (class, register, and patch) as image conditions.

### Gene preprocessing

Following best practices,^94,95^ we removed unwarranted genes (i.e., those absent from the official 10x Genomics panels) and transformed the gene expression matrix using the shifted natural logarithm. We avoided normalization because it can distort biologically meaningful variation in mRNA abundance across regions.^96^ For instance, tumor areas typically have higher count densities than stromal or immune regions. Applying global normalization may artificially inflate expression in regions with low content (e.g., immune niches), creating misleading patterns. Despite these steps, the data can remain highly noisy due to biological and technical batch effects.^97^ This may render the data manifold discontinuous and any generative model ill-conditioned. We hence applied k-nearest neighbor (k-NN) smoothing,^98^ which optionally aggregates information from neighboring cells or spots.

### Neural compression

Autoencoders^99^ (AEs) are neural networks designed to learn compact representations of data, primarily for tasks such as neural compression and feature extraction. They consist of two main parts: an encoder that maps the input data to a latent vector, and a decoder that reconstructs the original data from that vector (typically by minimizing the L2 loss). To prevent trivial identity mapping, the latent space $\mathcal{Z}$ is deliberately smaller than the input space $\mathcal{X}$, ensuring that the model learns informative features rather than memorizing the input. This encourages the network to find an efficient coding of the data, exploiting the assumption that high-dimensional observations lie on a lower-dimensional manifold. For Phoenix, we developed a new variant called the MLP-Mixer-AE. The original MLP-Mixer^20^ uses multilayer perceptrons^100^ (MLPs) to compute weighted linear combinations both within tokens (“channel mixing”) and across tokens (“sequence mixing”). By reducing or expanding the output dimension of the sequence mixer, we formed the encoder or decoder and thereby constructed an autoencoder. Phoenix employs an optional AE with an embedding dimension of 8 or 16.

### Generative modeling

Following the conditional flow matching (CFM) framework^101^, we constructed a family of conditional probability paths $(p_t(\cdot \mid x_0, x_1))_{t \in [0,1]}$ between source samples $x_0 \sim p_0$ and data samples $x_1 \sim p_1$. Marginalizing these conditional probability paths over the joint distribution $q(x_0, x_1)$ induces a time-continuous marginal probability path $(p_t)_{t\in[0,1]}$ that transports $p_0$ to $p_1$. The marginal probability path is generated by a marginal vector field $u_t(x)$. This aggregate field is intractable but can be expressed as the expectation of the simpler conditional vector fields $v_t(x \mid x_0, x_1)$, which govern the instantaneous motion of individual samples along the trajectories that define the conditional probability path (the density centered at that trajectory). Adopting a linear trajectory $x_t = (1-t)x_0 + t x_1$, the conditional vector field takes the closed form $v_t(x \mid x_0, x_1) = x_1 - x_0$. Therefore, it is feasible to train a neural network $u_\theta(t,x)$ to approximate these target fields via regression. Finally, integrating the time-dependent velocity field by solving the ODE $\dot{x} = u_\theta(t,x)$ computes the flow; this defines a deterministic map that pushes forward samples from the source distribution $p_0$ along the learned trajectories to the data distribution $p_1$. The time-evolution of these random samples constitutes the stochastic process $(x_t)_{t \in [0,1]}$. Implementation-wise, our neural network is a modified transformer^16^. We took the diffusion transformer^102^ and replaced it with modern components^18^ like Pre-Normalization^103^, RMSNorm^104^, and SwiGLU-MLP^105^. A cross-attention layer was added between each self-attention and feed-forward layer.^19^ The final model contains eight such blocks, each with an embedding dimension of 512, eight attention heads, and a head dimension of 64.

### Implementation details

Phoenix variants span all combinations of cohorts (CHUV, LMU, SNUH, TENX, TUM, UKER), gene panels (breast, colon, lung, skin, immuno-oncology, multi-tissue & cancer), spatial resolutions (100 µm, 55 µm, single-cell), and image resolutions (20×, 40×). Training was conducted on the JURECA pre-exascale modular supercomputer located at the Jülich Research Center, using distributed computation across multiple accelerators, totaling more than 10,000 GPU hours on nodes equipped with 4 x NVIDIA H100 NVL Tensor Core GPUs, each with 96 GB of HBM3 memory and 3.9 TB/s memory bandwidth. With a power consumption of 350–400 watts per device, the whole process consumed approximately 6,000 kWh, which, based on the average carbon intensity of the German energy grid (343 g CO₂e/kWh)^106^, corresponded to an estimated carbon footprint of about 2.0 tonnes CO₂e.

### Model inference

At inference time, we generate samples from our conditional flow matching model by numerically integrating the learned vector field. Given a conditioning signal $c$ and an initial sample $x_0 \sim \mathcal{N}(0, 1)$ drawn from the prior distribution, we obtain the target sample $x_1$ by solving the ordinary differential equation (ODE) $\frac{dx_t}{dt} = v_\theta(x_t, t, c)$ for $t \in [0, 1]$, where $v_\theta$ denotes the conditional velocity field parameterized by our neural network with learned parameters $\theta$. We integrate this ODE using the Dormand-Prince method (DOPRI5), an adaptive-step explicit Runge-Kutta solver of order 5(4) that adjusts its step size based on an embedded fourth-order error estimate. We set both the relative and absolute error tolerances to $1 \times 10^{-1}$ (i.e., $\mathrm{rtol} = \mathrm{atol} = 10^{-1}$), which we found to provide a favorable trade-off between sample quality and computational cost, allowing the adaptive solver to take relatively large steps in regions where the velocity field is smooth while refining the step size in more curved regions of the trajectory. Integration is performed over the unit interval $t \in [0, 1]$, with $x_0$ as the initial condition and the solution at $t = 1$ taken as the generated sample. The H&E-stained single-cell image used for conditioning can be obtained in two ways. The first follows a traditional computational pathology pipeline, in which the whole slide image is partitioned into 224 x 224 patches on a regular grid, and the model predicts the expression of the cell located closest to the center of each patch. The second leverages CellViT or HistoPlus to segment individual nuclei, after which a 224 x 224 patch is cropped around each segmented nucleus such that it is centered within the patch.

### Biological experiments

Single-cell atlases came from the Curated Cancer Cell Atlas (Weizmann Institute of Science) and Cell x Gene (Chan Zuckerberg Initiative). The cited studies provided marker genes and data for validation – as well as approaches, frameworks, and methods for downstream experiments. In addition, we used these tools for cell annotation: Tirosh method (scanpy), differential expression: Wilcoxon rank-sum test (scanpy); genomic analysis: copy number variants (infercnvpy); hypothesis testing: Student’s t-test (statsmodels), label transfer: ingest (scanpy); multiple testing: Benjamini–Hochberg procedure (statsmodels); spatial analysis: distances, neighbours, niches, proportions, locations (numpy, pandas, scipy, sklearn, squidpy); statistical learning: cross-validation, generalized estimating equations, linear mixed models, linear regression, logistic regression (sklearn); survival analysis: Cox proportional hazards, Kaplan-Meier estimator (lifelines, scikit-survival).

## Ethics statement

All research procedures were conducted in accordance with the Declaration of Helsinki. **CHUV:** All patients provided informed consent for the use of the tumor samples in this study. The protocol used for sample collection was approved by the local ethics committee (CER-VD, BASEC ID 2016-02094). **UKER:** All patients provided informed consent for the use of the tumor samples in this study. The protocol used for sample collection was approved by the local ethics committee and various protocol IDs (protocol code 4607, 2015; protocol code 97_18Bc, 2018; protocol code 22-343-B, 2022; and protocol code 217_18C, 2018). **LMU:** Sample collection and biological analysis of the cohorts provided to this study were approved by the local ethics committee, i.e., head and neck cancer data (reference no. 17-0116 and 23-0390) and sarcoma data (reference no. 20-824 and 23-0113).

## Data availability

The NCBI cohort is available at https://www.nature.com/articles/s41467-023-43458-x. The SNUH cohort is available at https://www.nature.com/articles/s41592-025-02770-8. The TENX cohort is available at https://www.10xgenomics.com/datasets. The BRCA cohort is available at https://www.modernpathology.org/article/S0893-3952(22)00349-0. The HGSOC cohort is available at https://www.cell.com/cell/fulltext/S0092-8674(23)00737-7. The HNSCC^107^ cohort is available at https://www.nature.com/articles/s41467-025-62386-6. The ovarian cohort is available at https://www.nature.com/articles/s41597-022-01127-6. The remaining cohorts can be requested from the respective data owners to bona fide researchers upon reasonable written request.

## Code availability

The model code is available at https://github.com/peng-lab/phoenix. The model weights are available at https://huggingface.co/peng-lab/phoenix. The spatial processing pipeline is available at https://github.com/peng-lab/spatialrefinery. The histology processing pipeline is available at https://github.com/peng-lab/HistoBistro. The spatial atlases and image patches / features are available at https://huggingface.co/datasets/peng-lab/phoenix.

## Acknowledgements

M.T. and R.H.G. are supported by the Helmholtz Association under the joint research school “Munich School for Data Science – MUDS”. A.M. received funding from the Bavarian Cancer Research Center (BZKF) to generate the Xenium sarcoma data and is supported by the Max-Eder Research Group Program of the German Cancer Aid (Deutsche Krebshilfe). L.M.B. is supported by the Else Kröner-Fresenius Foundation (EKFS) under the Immuno Oncology and Local Intervention (IOLIN) research program at the LMU University Hospital Munich. The study was supported by the German Cancer Consortium (DKTK) and DKTK Joint Funding, the DKFZ-MOST cooperation program (Ca-217), the Deutsche Forschungsgemeinschaft (SFB 1371 Project-ID 395357507 P12 to D.S.; DFG SA 1374/8-1 Project-ID 515991405 to D.S.; DFG SA 1374/7-1 Project-ID 515571394 to D.S., F.T.; DFG SA 1374/6-1 Project-ID 458890590 to D.S.); Deutsche Krebshilfe (#70117118 DEFEAT-PDAC of the German Pancreatic Cancer Alliance to D.S., F.T., C.F.; #70115743 to D.S.; #70116843 to D.S.).

## Author contributions

M.T. and R.H.G curated the dataset, developed the method, and ran the experiments. P.P. helped analyze the dataset. K.S. helped processing the dataset. G.P. helped conceptualize the study. T.K. and C.F. provided the pre-clinical analysis. C.W. provided the biological analysis. R.G. provided the clinical analysis. M.B. provided the COAD dataset. L.M.B., L.H.L., and T.K. generated the TMA dataset at LMU. L.B., P.J. and F.K. enabled and performed the LMU experiments. K.H. and R.G. provided the CHUV dataset. M.E. and C.M. provided the UKER dataset. A.M. provided the LMU dataset and supervised the biomedical experiments. D.S. provided the TUM dataset and supervised the biomedical experiments. F.J.T. supervised the machine learning experiments. T.P. supervised the entire study.

These authors contributed equally: Manuel Tran and Rushin H. Gindra.

## Corresponding authors

Correspondence to Andreas Mock, Fabian J. Theis, Dieter Saur, or Tingying Peng.

## Supplemental Tables

**Table 1:** Phoenix’s contributions relative to competing methods. Phoenix trains on a dataset one to two orders of magnitude larger than existing methods, covers more organs and genes, uniquely supports out-of-the-box inference, generalization to unseen tissues and cohorts, and in silico hypothesis testing, biomarker screening, and perturbation modeling. Checkmarks denote capabilities explicitly demonstrated in the original publication; crosses indicate those that were not. “Effective genes” refers to the number of genes modeled or predicted under the reported setting. HVG, highly variable genes; SVG, spatially variable genes.

| Contributions | Phoenix | BLEEP | DeepSpot | GHIST | SpatialEX |
| --- | --- | --- | --- | --- | --- |
| Spatial resolution | cell, spot | spot | cell, spot | cell, spot | cell, spot |
| Training dataset | 22.2M | < 1M | < 1M | < 1M | < 1M |
| Organs & tissues | 16 | 7 | 4 | 3 | 3 |
| Effective genes | 280-480 | 50 HVG | 100 HVG | 50 SVG | 280-379 |
| Out of the box predictions | ✓ | × | × | × | × |
| Generalization to unseen tissues | ✓ | × | × | × | × |
| Generalization to unseen cohorts | ✓ | × | × | × | × |
| In silico testing of hypotheses | ✓ | × | × | × | × |
| In silico screening of biomarkers | mKras<sup>G12D</sup>, tumor subtypes | × | × | × | × |
| In silico modeling of treatment | antibody, platinum therapy | × | × | × | × |

## Extended Figures

**Extended Data Fig. 1.**
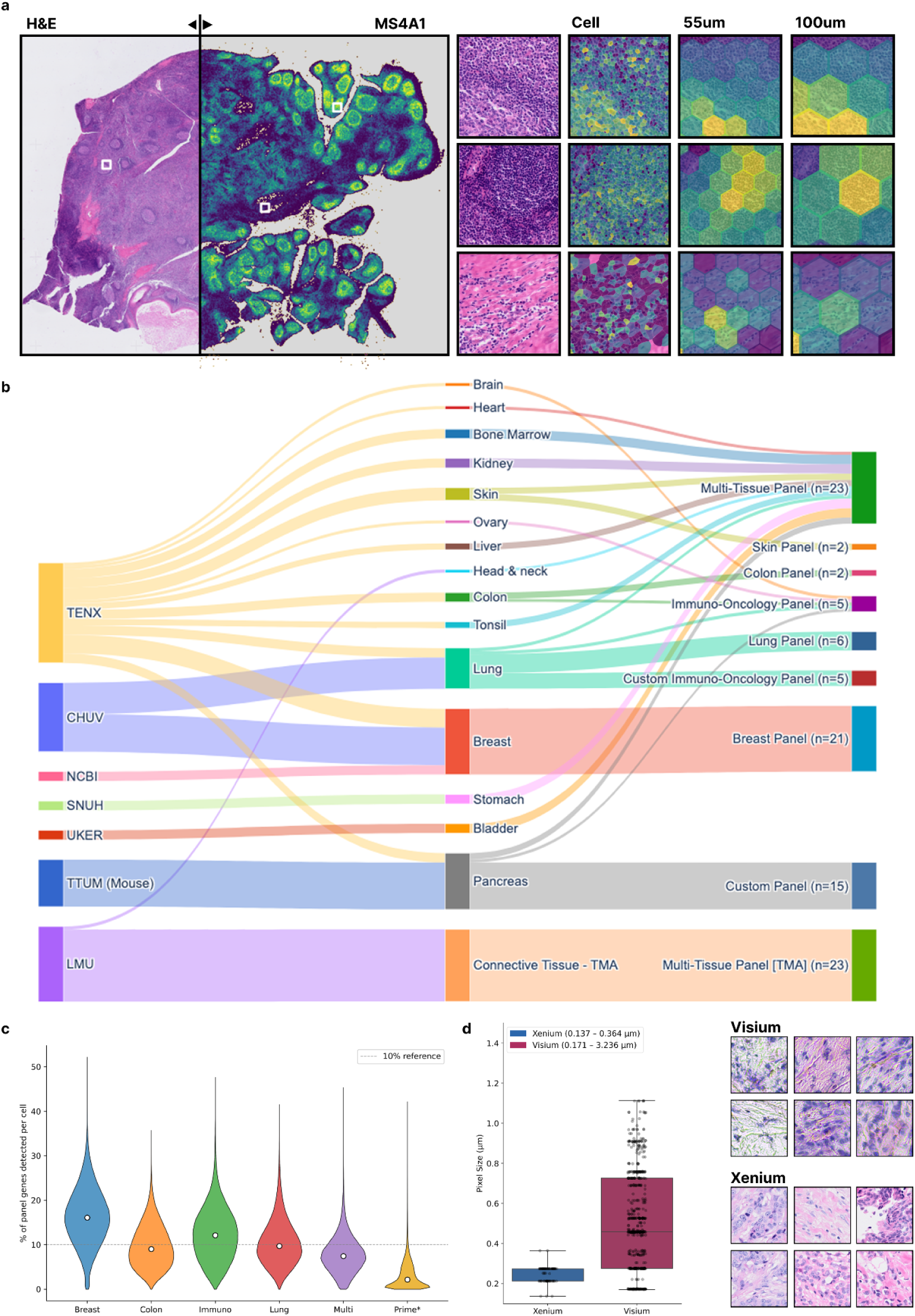
Dataset overview. **(a)** H&E-stained histology image of human tonsil reactive follicular hyperplasia profiled by Xenium In Situ spatial transcriptomics, with MS4A1 expression overlaid on the tissue (left). Example regions show MS4A1 expression at single-cell, 55 µm, and 100 µm resolutions (right). **(b)** The Nest dataset used for model training, along with mouse pancreatic ductal adenocarcinoma (mPDAC) and human sarcoma samples. **(c)** Percentage of panel genes detected per cell in each organ. **(d)** Image quality differs between Visium (CytAssist scanner) and Xenium (high-resolution scanner). Left: Smaller pixel size is better. Right: Visium tissue shows morphological distortions and artifacts absent in Xenium.

**Extended Data Fig. 2.**
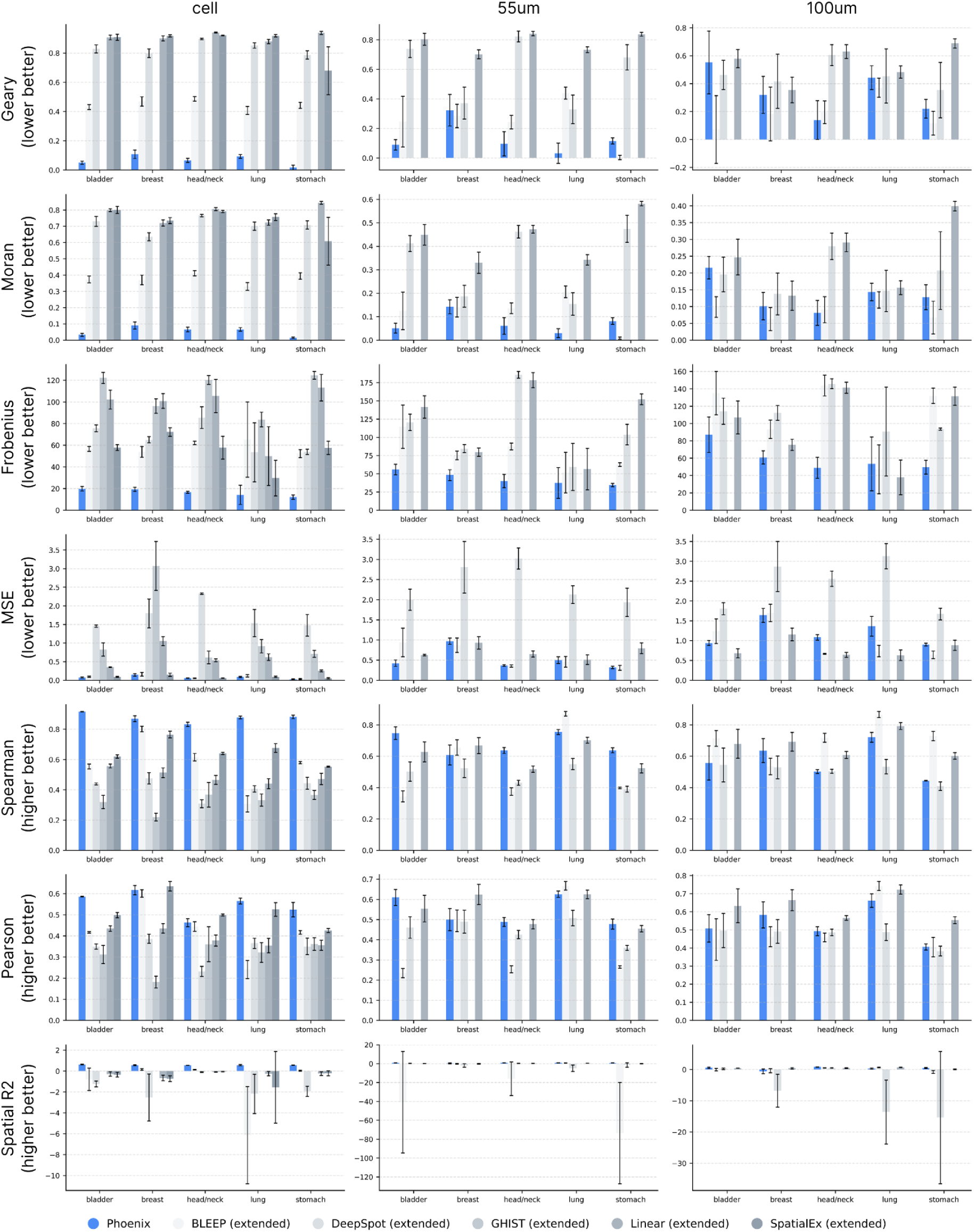
Benchmarking Phoenix against competing methods (BLEEP^24^, DeepSpot^25^, GHIST^26^, Linear^27^, SpatialEx^28^>) across bladder, breast, head & neck, lung, and stomach samples. Geary’s C looks at the differences between neighbors, making it more sensitive to local patterns (figure: 1 − C_true_ / C_pred_). Moran’s I compares each value to the global mean, making it better at detecting global patterns (figure: |C_true_ − C_pred_|). Frobenius measures the distance between two co-expression matrices, making it more effective at tracking regulatory networks (figure: ||cov_true_ − cov_pred_||_F_). Mean squared error (MSE), Spearman correlation, and Pearson correlation assess the continuous expression domain, treating sub-threshold values as structural zeros.^108–111^ Spatial R² extends the classical coefficient of determination to the spatial domain. It quantifies the proportion of spatial autocorrelation present in the ground-truth observations that is reproduced by the model predictions.

**Extended Data Fig. 3.**
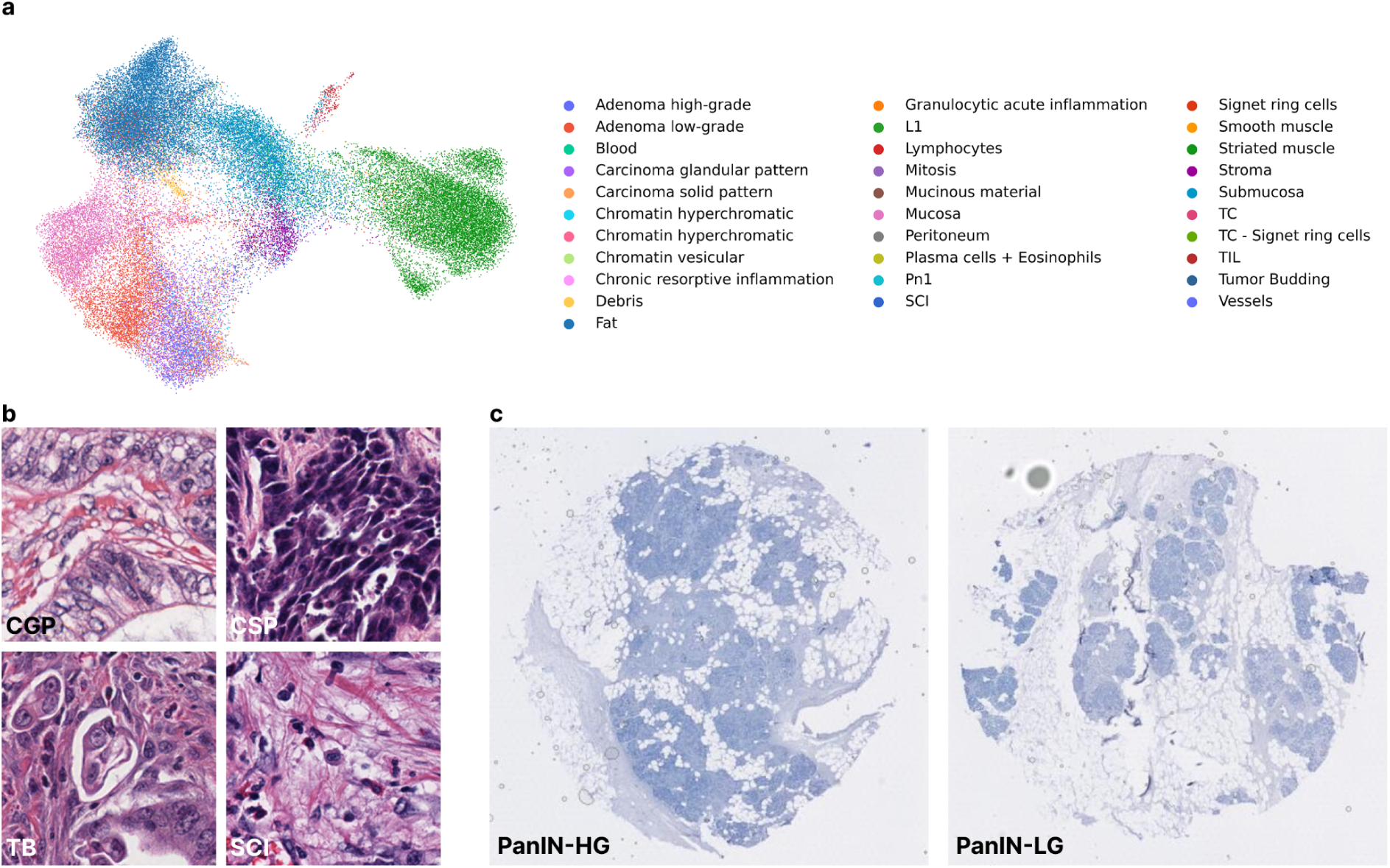
Datasets underlying Phoenix-driven discoveries. **(a)** UMAP of predicted gene expression derived from the colorectal cancer (CRC) dataset used in Fig. 3d–f, with each cell annotated by a pathologist. **(b)** Representative H&E-stained histology patches of colorectal adenocarcinoma showing glandular growth pattern (CGP), solid growth pattern (CSP), tumor budding (TB), and single-cell invasion (SCI). **(c)** Representative H&E-stained histology slides of pancreatic intraepithelial neoplasia (PanIN), showing high-grade (PanIN-HG; left) and low-grade (PanIN-LG; right) from the dataset used in Fig. 3g–i.

**Extended Data Fig. 4.**
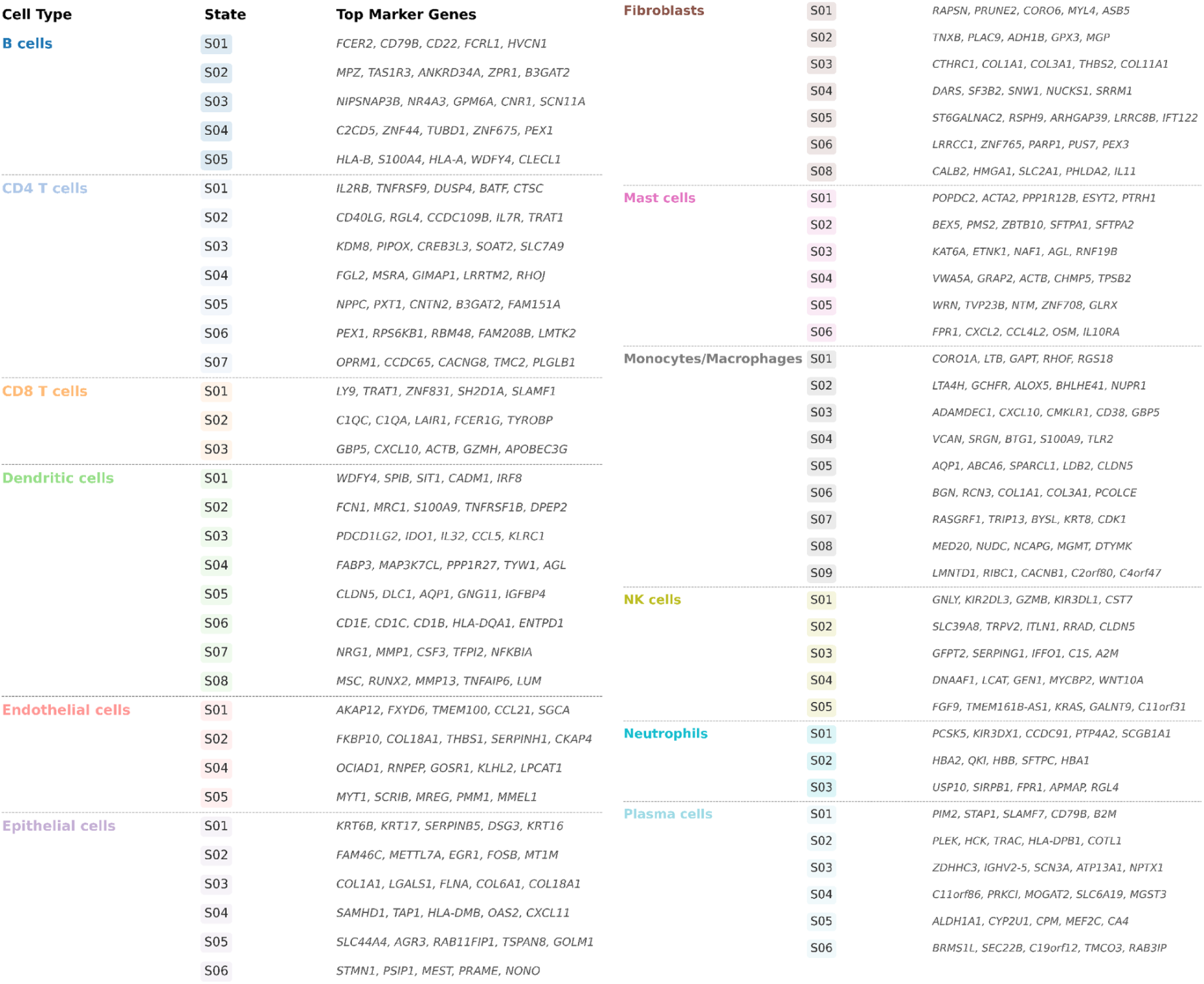
Key marker genes defining different immune, endothelial, epithelial, and fibroblast cell states from the two original Ecotyper studies.^54,55^ Cell states are classified by marker gene expression.

**Extended Data Fig. 5.**
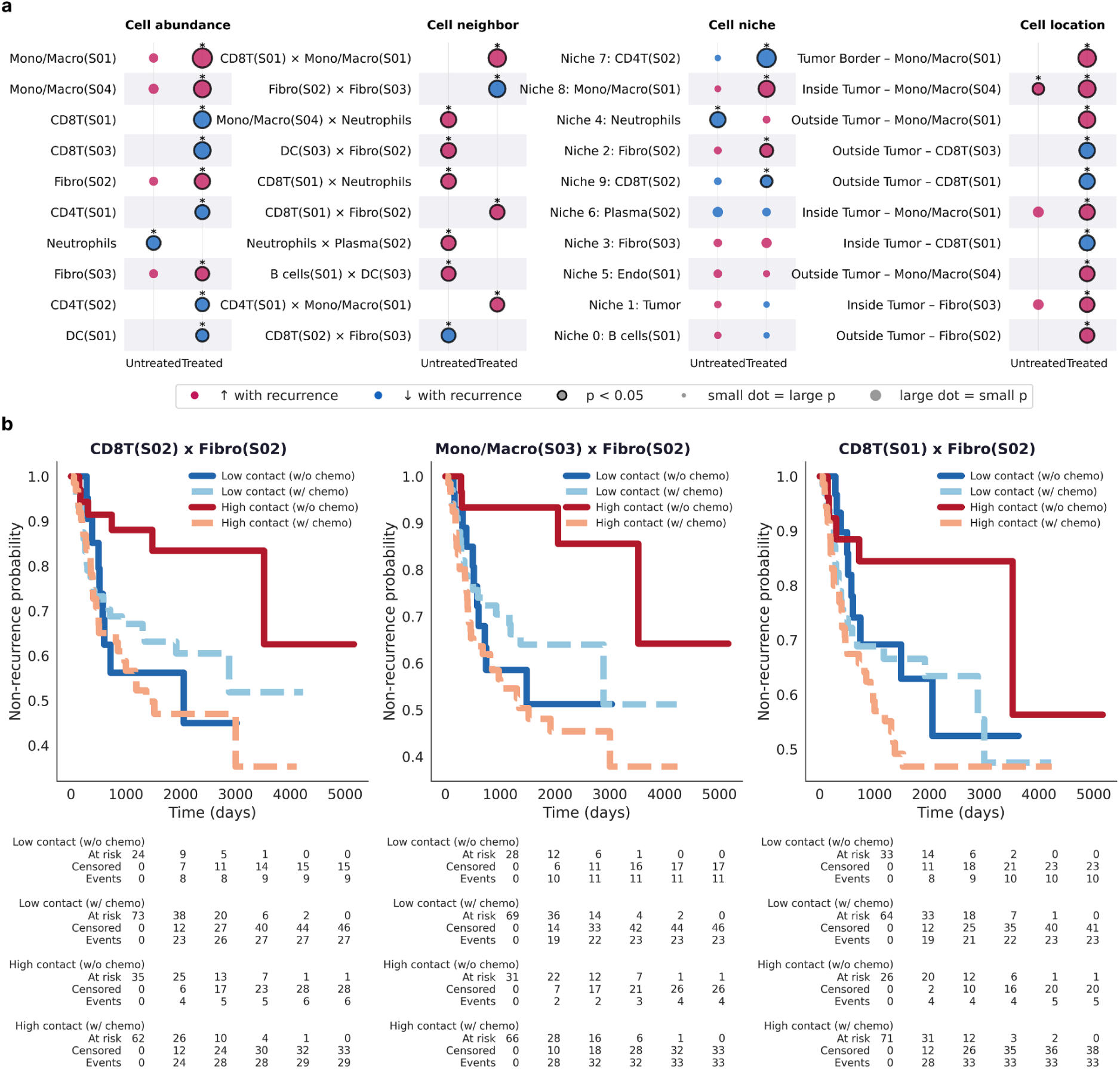
Supporting data for drug response experiments. **(a)** Cell abundance, neighbors, niches, and locations in head and neck squamous cell carcinoma (HNSCC) patients, stratified by recurrence and treatment (surgery alone vs. surgery + adjuvant radiotherapy / radiochemotherapy). **(b)** Kaplan–Meier recurrence estimates in HNSCC, stratified by chemotherapy and the most robust (top-3 selected) predictive spatial biomarkers: CD8T(S02) × Fibro(S02), Mono/Macro(S03) × Fibro(S02), and CD8T(S01) × Fibro(S02).

**Extended Data Fig. 6.**
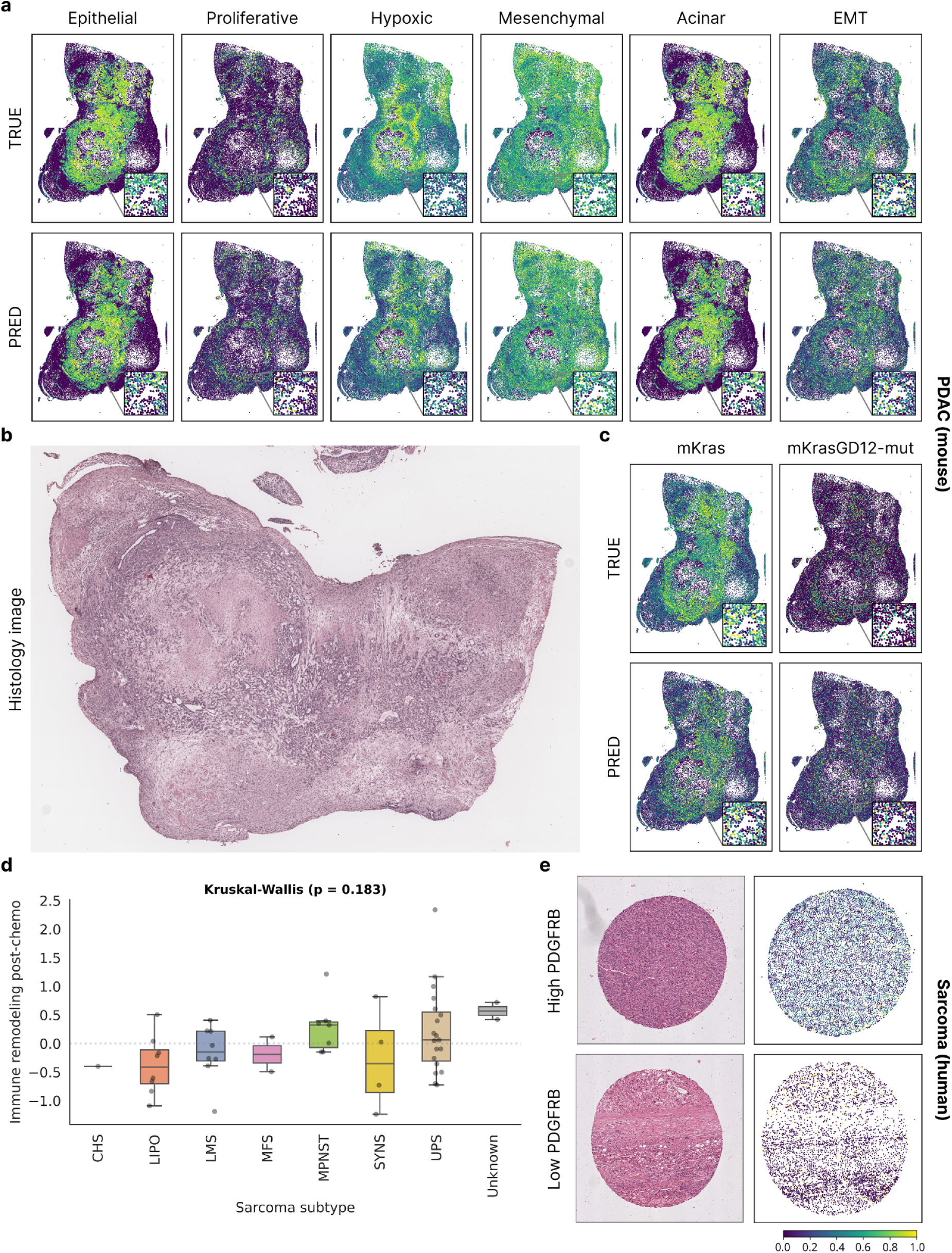
Supporting data for mesenchymal cancer and mouse model experiments. **(a–c)** Mouse PDAC. **(a)** Spatial expression of mPDAC subtype markers: epithelial (Cldn18, Krt19), proliferative (Mki67, Top2a), hypoxic (Hif1a, Vegfa), mesenchymal (Fn1, Thy1), acinar (Cdh1, Krt19), and EMT (Cdh2, Zeb1). **(b)** H&E-stained histology image for the sample in (a). **(c)** Spatial expression of mKras and mKrasG12D-mut. **(d–e)** Human sarcoma. **(d)** Sarcoma subtype, as classified by a pathologist using OncoTree codes, does not predict immune remodeling after chemotherapy (Kruskal-Wallis, p = 0.183). **(e)** H&E-stained histology image and experimental expression of PDGFRB in a core with high (top) and low (bottom) expression.

**Extended Data Fig. 7.**
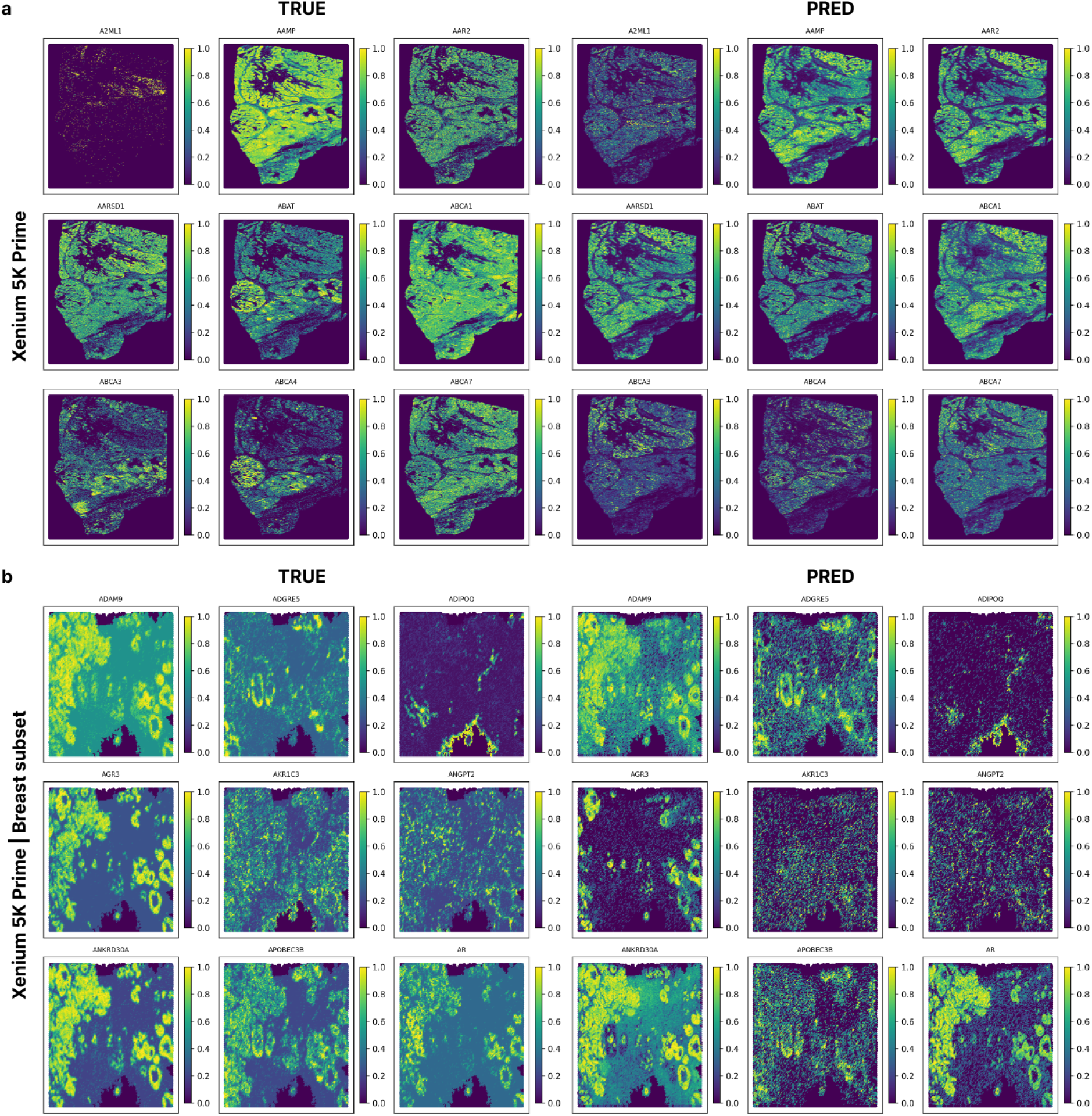
Phoenix trained on Xenium 5K Prime data generalizes poorly to new samples. **(a)** A Phoenix model trained on mixed-tissue Xenium 5K Prime samples and evaluated on a held-out ovarian Xenium 5K Prime sample. **(b)** A Phoenix model trained on mixed-tissue Xenium 5K Prime samples (breast panel genes only) and evaluated on a held-out Xenium breast panel sample. In both panels, ground truth expression (left) is shown alongside predicted expression (right). Predictions consistently underestimate expression levels, indicating that low sensitivity is the primary limitation when generalizing across samples on this platform.

